# Nuclear-cytoplasmic compartmentalization of the herpes simplex virus 1 infected cell transcriptome is co-ordinated by the viral endoribonuclease vhs and cofactors to facilitate the translation of late proteins

**DOI:** 10.1101/415497

**Authors:** Kathleen Pheasant, Carla Moller-Levet, Juliet Jones, Daniel Depledge, Judith Breuer, Gillian Elliott

## Abstract

HSV1 encodes an endoribonuclease termed **<u>v</u>**irion **<u>h</u>**ost **<u>s</u>**hutoff (vhs) that is produced late in infection and packaged into virions. Paradoxically, vhs is active against not only host but also virus transcripts, and is involved in host shutoff and the temporal expression of the virus transcriptome. Two other virus proteins - VP22 and VP16 – are proposed to regulate vhs to prevent uncontrolled and lethal mRNA degradation but their mechanism of action is unknown. We have performed dual transcriptomic analysis and single-cell mRNA FISH of human fibroblasts, a cell type where in the absence of VP22, HSV1 infection results in extreme translational shutoff. In Wt infection, host mRNAs exhibited a wide range of susceptibility to vhs ranging from resistance to 1000-fold reduction, a variation that was independent of their relative abundance or transcription rate. However, vhs endoribonuclease activity was not found to be overactive against any of the cell transcriptome in Δ22-infected cells but rather was delayed, while its activity against the virus transcriptome and in particular late mRNA was minimally enhanced. Intriguingly, immediate-early and early transcripts exhibited vhs-dependent nuclear retention later in Wt infection but late transcripts were cytoplasmic. However, in the absence of VP22, not only early but also late transcripts were retained in the nucleus, a characteristic that extended to cellular transcripts that were not efficiently degraded by vhs. Moreover, the ability of VP22 to bind VP16 enhanced but was not fundamental to the rescue of vhs-induced nuclear retention of late transcripts. Hence, translational shutoff in HSV1 infection is primarily a result of vhs-induced nuclear retention and not degradation of infected cell mRNA. We have therefore revealed a new mechanism whereby vhs and its co-factors including VP22 elicit a temporal and spatial regulation of the infected cell transcriptome, thus co-ordinating efficient late protein production.

**Author Summary:** Herpesviruses are large DNA viruses that replicate in the nucleus and express their genes by exploiting host cell mRNA biogenesis mechanisms including transcription, nuclear export, translation and turnover. As such, these viruses express multiple factors that enable the appropriation of cellular pathways for optimal virus production, and work in concert to shut off host gene expression and to overexpress virus genes in a well-described cascade that occurs in a temporal pattern of immediate-early, early and late proteins. We have analysed global and single cell changes in the host and virus transcriptome to uncover a novel mechanism by which the viral endoribonuclease, termed vhs, turns off early virus gene expression. This is achieved through the vhs-induced nuclear retention of the entire infected cell transcriptome at the onset of late gene expression. To enable the switch from early to late protein production the virus then requires a second factor called VP22 to specifically inhibit the nuclear retention of late transcripts allowing their translation in the cytoplasm. In this way, HSV1 elicits a temporal and spatial regulation of the infected cell transcriptome to co-ordinate efficient late protein production, a process that may be relevant to herpesviruses in general.

## Introduction

Herpesviruses exhibit two major characteristics of gene expression during lytic infection: a global shutoff of host gene expression, and a temporal pattern of virus gene expression resulting in a cascade of immediate-early (IE), early (E) and late (L) protein synthesis, such that L genes encoding the virus structural proteins are expressed optimally after DNA replication [1]. These two features are interlinked through the complex activities of a number of virus factors which regulate and usurp cellular post-transcriptional RNA biogenesis steps, including splicing, nuclear export, stability and association with the translation machinery. To date, the best-characterised protein shown to be involved in RNA biogenesis is the herpes simplex virus 1 (HSV1) IE protein ICP27 (and homologues) which is involved in the shutoff of host translation [2, 3] by inhibiting the export of spliced cellular mRNAs [4], and the expression of virus proteins, being essential for the export of unspliced L viral transcripts in to the cytoplasm for their translation [2, 5-7]. Several herpesviruses encode their own endoribonucleases factors that also regulate host shutoff and the kinetics of virus gene expression, such as the HSV1 vhs (**<u>v</u>**irion **<u>h</u>**ost **<u>s</u>**hutoff) [8-11] and the KSHV sox (**<u>s</u>**hut**<u>o</u>**ff and **<u>e</u>**xonuclease) [12] proteins. These factors degrade host mRNAs in a global fashion, an activity that is believed to be important for counteracting host cell responses to virus infection [13], with multiple mRNAs for antiviral proteins degraded during infection [14-19]. vhs degrades mRNAs by binding to the cellular translation initiation machinery through the eIF4F cap-binding complex and cleaving the bound transcripts [20-23] implying a potential lack of discrimination between cellular and viral transcripts. Indeed, although non-essential in tissue culture, the fact that IE mRNAs are present at higher levels in cells infected with a Δvhs virus than in Wt infected cells [13], suggests that vhs is involved in regulating the temporal transition from IE to E gene expression by actively degrading IE mRNAs [24, 25]. Moreover, because vhs is packaged into the tegument of the virion [9, 26], it has the capacity to act at two stages of infection – very early in infection after incoming vhs has been delivered to the cytoplasm [27, 28], and at later times when it is newly synthesized, a time when significant global reduction in host cell mRNAs is readily detectable [29].

Given the apparent lack of selectivity for cellular mRNAs over viral mRNAs by vhs [30], it is considered that the high levels of vhs protein produced at later times of infection would be detrimental to virus infection leading to eventual total shutoff of virus protein synthesis. Two other virus proteins – VP22 and VP16 – are expressed around the same time as vhs and form a trimeric complex with it [31-33], leading to the proposal that they neutralise the RNase activity of vhs. In the absence of either VP22 or VP16, vhs would therefore ultimately degrade virus mRNA in an unrestrained fashion leading to complete shutoff of virus protein synthesis. In agreement with this model, deletion of VP16 is lethal to the virus causing complete translational shutoff at intermediate and late times of infection and a block to virus replication [34, 35]. However, in the case of VP22, although it has been reported by some that deletion of this gene is lethal to the virus in the presence of functional vhs [36], we and others have generated replication-competent, vhs-positive, VP22 deletion viruses which replicate with minimal defect in Vero cells [37-40]. Hence, the interplay of these proteins in the regulation of gene expression remains unclear.

Although vhs-induced translational shutoff is generally considered to be through its endoribonuclease cleavage of cytoplasmic mRNAs followed by Xrn1 exonuclease degradation [41], we have recently published that transient expression of vhs results in the nuclear retention of not only its own but co-expressed mRNAs, in a negative feedback loop that results in shutdown of translation of these transcripts in the expressing cell [42]. This newly-defined nuclear retention results in translational shutoff but is mechanistically different to the model of simple degradation of mRNAs located on ribosomes. However, together with the associated relocalisation of polyA binding protein (PABP) to the nucleus these results are consistent with studies reported for transiently expressed KSHV sox protein [43]. Moreover, the observation that this vhs-induced nuclear retention was overcome during transient transfection by co-expression of VP16 and VP22 pointed to a novel effect of these proteins on the translational shutoff activity of vhs [42]. Given these results, we have now conducted a comprehensive analysis of the role of VP22 in vhs activity during infection. We have identified human fibroblast cells (HFFF) as a cell type in which - unlike Vero cells – our Δ22 virus exhibits extreme translational shutoff, and have used dual transcriptomics and mRNA FISH to investigate global and single cell changes in the cell and virus transcriptome in the presence and absence of VP22. Despite complete translational shutoff, vhs activity against cell transcripts was not enhanced but rather was delayed when VP22 was not present. Moreover, the total virus transcriptome was only modestly reduced in the absence of VP22, but this reduction was specific to L transcripts. We have identified a novel role for vhs in the nuclear retention of all virus transcripts in the nucleus, with IE and E transcripts exhibiting nuclear retention at a time that correlates with the onset of vhs expression. Intriguingly, VP22 was required to overcome the nuclear retention activity of vhs on L transcripts, while a variant of VP16 which cannot interact with VP22 exhibited a phenotype intermediate between Wt and Δ22, suggesting that VP16 enhances the role of VP22 in regulating the compartmentalisation of the infected cell transcriptome. Hence, the characteristic translational shutoff seen in Δ22 infected cells is the consequence of dysregulated mRNA export rather than degradation. These results not only unveil a new mechanism for regulating the nuclear export of mRNAs for L protein expression in HSV1 infection, but also identify specific roles for vhs and VP22 in co-ordinating the specificity of this retention and export.

## Results

### Translational shutoff in HSV1 infected cells

To measure the contribution of the virus factors VP22 and vhs to translation in HSV1 infected cells, we carried out metabolic labelling of a range of cells infected with Wt (strain 17), Δ22 or Δvhs viruses. We have previously shown that our Δ22 virus replicates efficiently in Vero cells [37, 38], so this cell type was used as a reference together with a range of human cell types including HeLa, HaCaT or HFFF cells. The results indicated that after labelling for 1 hour at 15 hours after infection, the level of translation was broadly similar in Wt and Δ22 infected Vero cells, but a degree of translational shutoff was apparent in the Δ22 infections of all the human cells tested (Fig 1A). Most notably, almost complete shutoff occurred in the primary human fibroblast cell-type HFFF (Fig 1A). By contrast, Δvhs infected cells exhibited similar labelling levels to Wt infection in all cell-types although the profiles differed slightly (Fig 1A). To determine the kinetics of translation shutoff during Δ22 infection of HFFF, metabolic labelling was carried out at different times after infection (Fig 1B). This revealed that at 5 h, the labelling profile of all three viruses was similar and comparable to uninfected cells, indicating that the virus had not yet taken over the translation machinery of the cell at this time (Fig 1B, 5 hpi). By 10 h, the majority of proteins translated in both the Wt and Δvhs infections were viral proteins, although the Δvhs infection contained a stronger background of alternative presumably cellular proteins (Fig 1B, 10 hpi). By contrast, translation in the Δ22 infected cells had already begun to shut down at this time, and by 15h was almost completely halted in comparison to either Wt or Δvhs ((Fig 1B, 15 hpi). The relative ability of these three viruses to form plaques on Vero or HFFF cells also reflected this degree of translational shutoff, with Δ22 unable to plaque on HFFF cells but showing only a ∼40% reduction in plaque size on Vero cells (Fig 1C) [38]. Moreover, infection with viruses lacking the virion kinase UL13, or the neurovirulence factor ICP34.5 – virus proteins that have been shown to have a role in translation in HSV1 infected cells [44-46]– resulted in neither a global reduction in translation, nor an extreme decrease in plaque size in HFFF (S1 Fig). This confirmed the direct importance of VP22 in the shutoff phenotype in HFFF.

**Fig 1.**
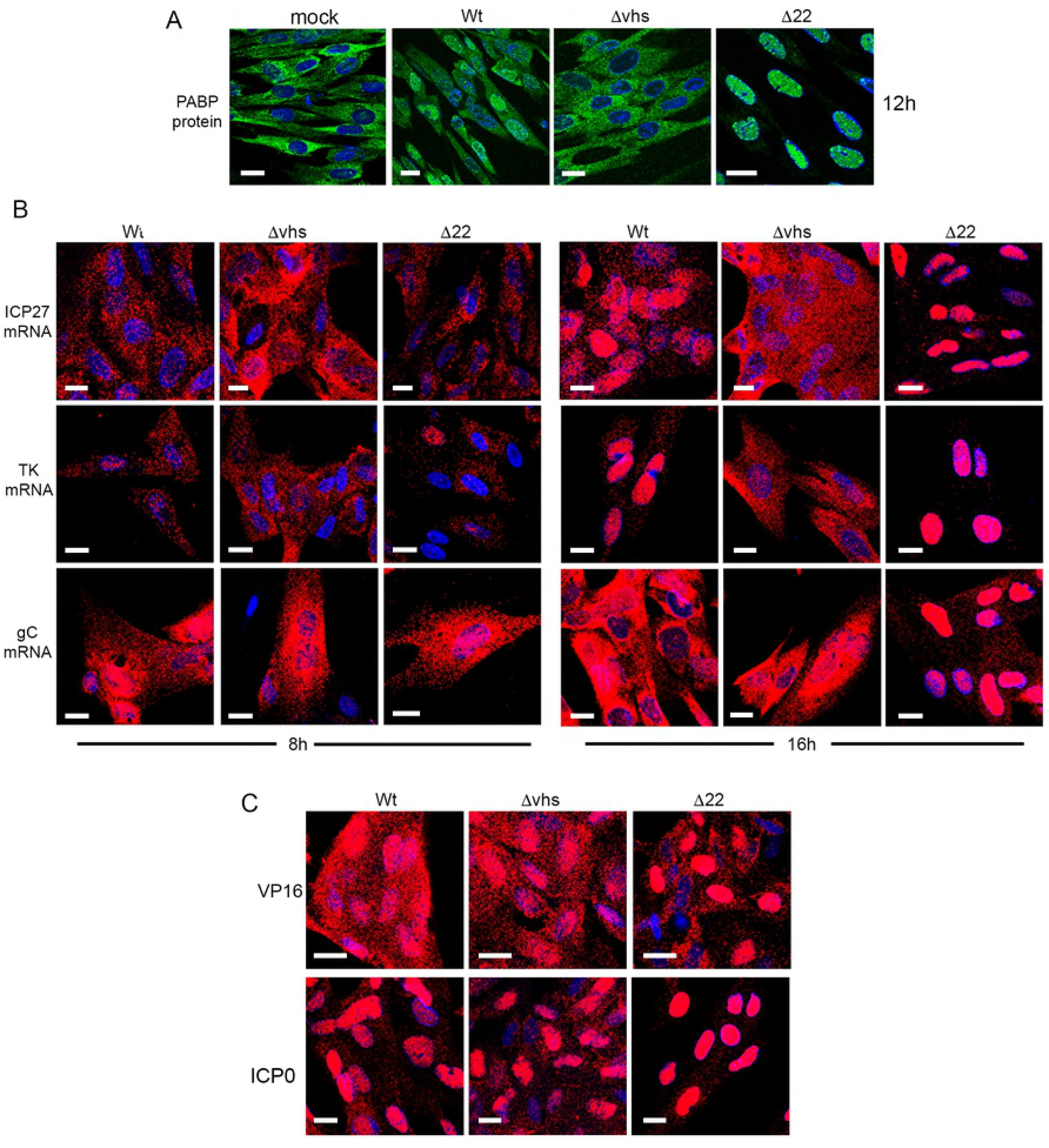
Translational shutoff during HSV1 infection in the absence of VP22 expression. (A).The indicated cell types were infected with Wt (s17), Δ22 or Δvhs viruses at a multiplicity of 2, and 16 hours later were incubated in the presence of [35S]-methionine for a further 60 mins. The cells were then lysed and analysed by SDS-PAGE followed by autoradiography.(B) HFFF cells were treated as in A, and metabolic labelling with [35S]-methionine was carried out at the indicated times after infection. (C) Confluent monolayers of HFFF were infected with ∼ 30 plaque forming units of each virus and plaques allowed to develop for 5 days before fixing and staining with crystal violet. (D) HFFF cells infected as in A were harvested at 16 hours and analysed by SDS-PAGE and Western blotting with antibodies as indicated. (E) Vero cells were left untreated, or treated for 20 hours with 1000 units/ml of interferon β, prior to the titration of the indicated viruses. The mean and ± standard error of the data is given from three independent experiments. Statistical analysis was carried out using an unpaired, two-way student’s t test. ns, p > 0.05. *** p < 0.001. (F) The samples from D were analysed by SDS-PAGE and Western blotting with the indicated antibodies.(G) HFFF cells infected with Wt, Δ22 or Δvhs viruses at a multiplicity of 2 were harvested at the indicated times after infection (in hours) and analysed by Western blotting for VP16,α tubulin and phospho-eIF2α.

The metabolic labelling profile and timing in HFFF also correlated with detection levels of individual virus proteins by Western blotting where a number of virus proteins – in particular the L proteins and specifically glycoproteins tested - were greatly reduced in HFFF cells infected with the Δ22 virus compared to either Wt or Δvhs infected cells at 16h (Fig 1D). The IE and E proteins that were examined (ICP27 and TK in Fig 1D) were however present at a similar level in Δ22 and Wt infection presumably reflecting their translation prior to the onset of shutoff. Moreover, as shown by others [13, 25], the IE and E proteins were overexpressed in Δvhs infected HFFF cells (Fig 1D), indicative of failure of vhs to degrade these transcripts. Of note, and as we have reported recently in HeLa cells, vhs was itself poorly translated in Δ22 infected HFFF cells [42, 47].

Unlike HFFF cells, Vero cells are unable to produce interferon but are fully responsive to it [48]. To understand why the Δ22 virus exhibited such extreme translational shutoff in HFFF, we first determined if the Δ22 virus might be more sensitive to the actions of interferon by performing a plaque reduction assay, whereby the virus was titrated on Vero cells that had been left untreated or pre-treated with recombinant interferon β. The effect of interferon on the titre of the Δ22 and Δvhs viruses was compared to the Wt s17, which is known to be generally resistant to its activity [49], and a s17 derived virus lacking ICP34.5 (Δ34.5) which is highly sensitive to interferon [50] (Fig 1E). In this case, both the Δ22 and Δvhs viruses were judged to be relatively resistant to the actions of interferon β and their titres were reduced by no more than 30-fold compared to 10-fold for Wt virus (Fig 1E). By contrast, the Δ34.5 virus showed over 2000-fold reduction (Fig 1E). In addition, we demonstrated that there was no induction of IRF3 phosphorylation in Δ22 infection that might indicate increased sensing of virus infection in these cells (Fig 1F), whereas cells infected with the Δvhs virus contained enhanced phospho-IRF3 levels as expected for its known role in targeting antiviral responses [13]. Finally, we also examined the Δ22-infected cells for the presence of hyperphosphorylated eIF2α that could indicate host-induced shutoff of translation. Uninfected HFFF already contained relatively high levels of phospho-eIF2α, and as expected from earlier studies [13, 51, 52], this was reduced in cells infected with Wt but not.Δvhs HSV1 (Fig 1F). Nonetheless, in spite of the extreme translational shutoff in Δ22 infected cells, phosphorylation of eIF2α was not upregulated in the absence of VP22 (Fig 1F). Analysis of a time course of infection for all three viruses showed an immediate reduction in eIF2α phosphorylation in the first 2 hours of infection, but that in Δvhs-infected cells, phosphorylation recovered quickly to uninfected cell levels by 4 hours suggesting that vhs is required to maintain the reduction in eIF2α phosphorylation (Fig 1G). In all infections, further reduction of eIF2α phosphorylation correlated with the start of L protein synthesis from 8h onwards (Fig 1G), a feature that would be consistent with the activity of ICP34.5 [53]. In short, these data indicate that the translational shutoff seen during Δ22 infection was not a consequence of cell-induced inhibition of the translation machinery. It is therefore noteworthy that as we have seen before [54], the Δ22 virus does not have the phenotype of a Δvhs virus, despite the low level of vhs expressed in the absence of VP22 (Fig 1D) [37, 42].

### Transcriptomic analysis reveals highly variable susceptibility to vhs activity in HSV1 infected cells

An obvious reason for translational shutoff is the reduction of the pool of mRNA available for translation. As such, the current model predicts that vhs endonuclease activity is unregulated in HSV1 infected cells where VP22 is absent and hence cellular and viral mRNAs would be predicted to be hyper-degraded [36]. To comprehensively measure the extent of vhs activity in HSV1 infected HFFF cells and compare the relative activity of vhs in the presence and absence of VP22 on both cellular and viral transcripts, dual transcriptomic analysis of Wt (strain 17) and Δ22 infected HFFF was carried out at 0, 4 and 12 hours after infection. RNAseq was performed on 5 biological replicates for each condition, with sequence reads being mapped to both the HSV1 genome and the human genome, and normalisation and filtering carried out as described in Methods. Over 11,000 cellular transcripts were detected in the HFFF libraries prepared from uninfected cells (as indicated in Tables S1 to S6). Mapping the reads obtained for each library to the human and HSV-1 transcriptome indicated that the library composition comprised on average only 3% virus reads at 4 h, but by 12 h this had risen to close to 75% (Fig 2A). This is quite different to the situation previously shown for VZV where the virus has been shown to make up around 20% of the transcriptome [55], and reflects the massive impact that HSV1 has on the total cell transcriptome. Normalised and averaged library counts of each gene were used to determine the differential expression of each detected transcript at 4 h and 12 h in Wt infected cells in comparison to uninfected cells, expressed as Log_2_ fold change (FC) to uninfected (S1 & S2 Tables). Scatter plots showing average counts per million of transcripts of uninfected versus 4 h or 12 h libraries revealed that there were only small differences in abundance of most cellular transcripts at 4 h, but that by 12 h the vast majority of cell transcripts were reduced in abundance in accordance with vhs activity (Fig 2B). Around 100 cellular transcripts were upregulated > Log_2_ FC of 1 as early as 4 h, although the majority of the obviously increased transcripts were viral in origin (Fig 2B, 4h, cell transcripts in black; virus transcripts in green). Three quarters of the upregulated cellular transcripts were identified as representing interferon stimulated genes (ISGs) by screening those transcripts that were increased against the interferome database (http://www.interferome.org) eg IFITs 1 and 2 as identified in Fig 2B (see also S2 Fig and S1 Table). By 12 h, 98% of cellular transcripts were reduced more than 2-fold in Wt infection compared to uninfected cells, a result taken to reflect the activity of the vhs endoribonuclease in the cell (Fig 2B, 12h; S2 Table).

**Fig 2.**
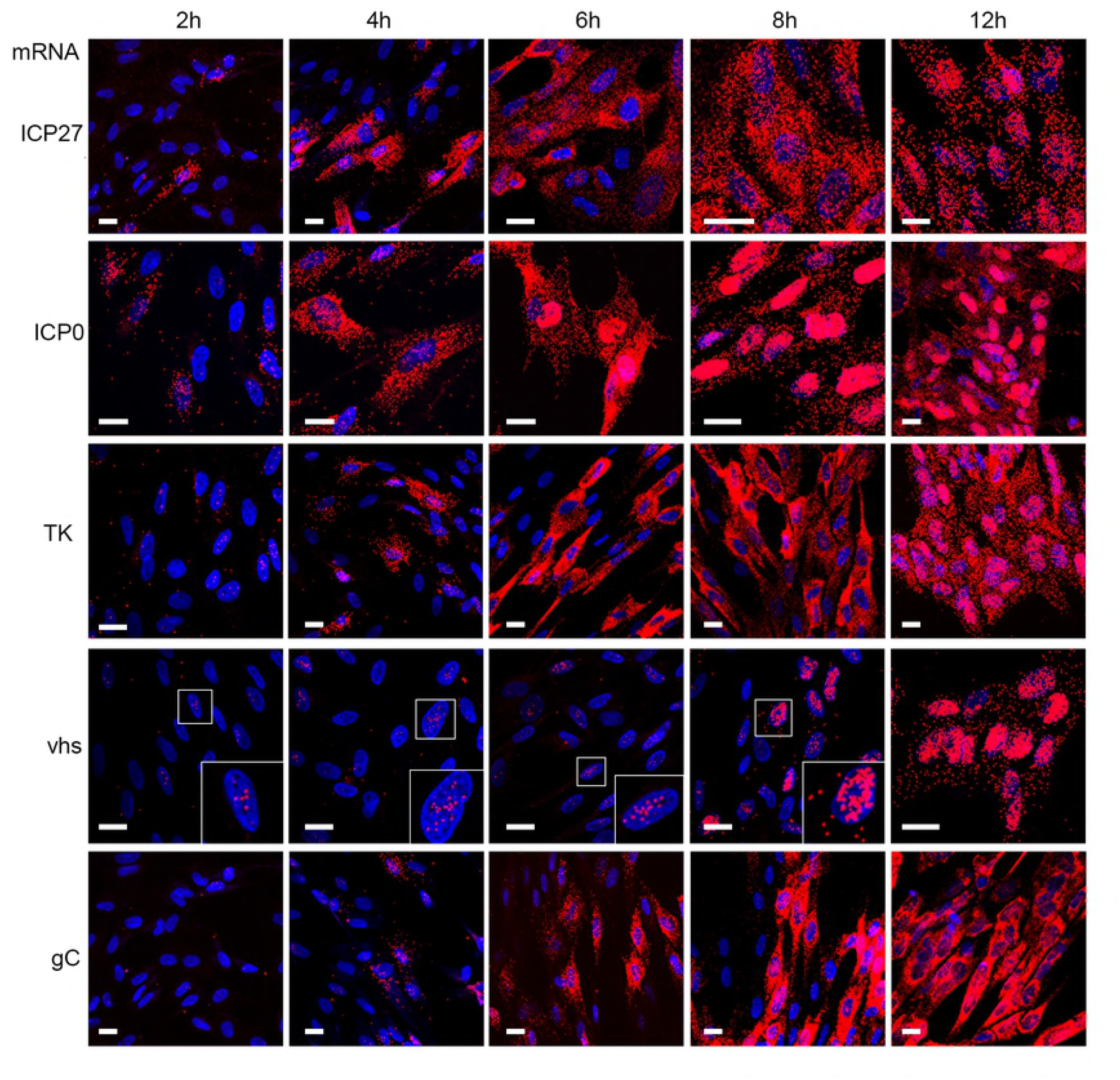
Dual transcriptomic analysis of human fibroblast cells infected with HSV1. HFFF cells were left uninfected, or infected with HSV1 (s17) at a multiplicity of 2. At 4 or 12 h.p.i., total RNA was purified and used for library preparation followed by sequencing. A total of 5 biological replicates were sequenced for each condition. (A) The proportion of reads mapped in each condition to either the human (blue) or HSV1 (red) transcriptome. (B) Differential expression analysis of cell and virus transcripts was conducted using EdgeR as described in Methods. Differences in the number of reads mapped to cell (black circles) and virus (green circles) transcripts were plotted as scatter plots comparing results at 4 and 12 hours to uninfected cells. The 12 h results are also represented in a volcano plot indicating the high level of significance for the detected changes (right hand panel). (C) The reads obtained for the virus transcriptome were mapped to the virus genome for 4 (red) and 12 (green) hours. Numbers in parentheses represent maximum read counts per million obtained in each condition. The location of the TK, ICP27 and gD genes are indicated by arrows. (D) HFFF cells grown in slide chambers were infected with HSV1 (s17) at a multiplicity of 2 fixed at 4 or 12 h, and subjected to multiplex mRNA FISH with probes to genes representing IE (ICP27 in cyan), E (TK in red) and late (gD in green) transcripts. Nuclei were counterstained with DAPI (blue). Scale bar = 20 μm.

Mapping of the virus reads across the HSV1 genome at both time points indicated that at 4 h, as expected, the reads mapped predominantly to IE genes, such as UL54 (ICP27) and US1 (ICP22), or E genes such as UL23 (TK), UL29 and UL39 (Fig 2C, 4h). However, by 12 h the predominant transcription units covered the L genes including UL19 (major capsid protein), UL48 and UL49 (tegument proteins VP16 and VP22), UL27 and UL44 (glycoproteins gB and gC), and across the entire Us region of the genome (Fig 2C, 12h). The relative transcription of representative IE (ICP27), E (TK) and L gene (gD) transcripts was further confirmed at the single cell level using multiplex mRNA FISH which indicated that all cells in the population contained similar levels of each virus transcript, with ICP27 and TK levels changing little from 4h to 12 h, but gD increasing significantly, in line with the transcriptomic results (Fig 2D). Such single-cell studies of mRNA provide scope not only for analysing relative levels but also relative localisation of individual transcripts.

Further analysis of the ∼ 11,000 cellular transcripts revealed that there was a vast difference in the relative differential expression of these transcripts at 12 h, with some transcripts reduced by as much as Log_2_ FC of −10, while others were hardly altered (Fig 2B, 12h; S2 Table). To confirm that this variability detected by RNAseq truly reflected the relative abundance of transcripts in the RNA samples being analysed rather than mapping artefacts, we carried out qRT-PCR analysis on two independent RNA samples that had been prepared in the same way as the RNA for the RNAseq libraries. The relative levels of transcripts representative of upregulated, unaltered, and those exhibiting a range of susceptibility during infection broadly validated the RNAseq data (S3 Fig). To further confirm that the change in these transcripts was specific to the activity of vhs, qRT-PCR was carried out on RNA from HFFF cells infected with Wt or Δvhs virus at a multiplicity of 2 and harvested 16 h after infection. The reduction of all transcripts tested – those exhibiting both high and low susceptibility to degradation – was shown to be dependent on the presence of vhs during infection (Fig. 3A). Furthermore, three representative ISGs (IFIT1, IFIT2 and Herc5) were shown to be greatly induced at 16h in Δvhs compared to Wt infected cells (Fig 3B), confirming the role that vhs plays in degrading these induced antiviral transcripts [13].

**Fig 3.**
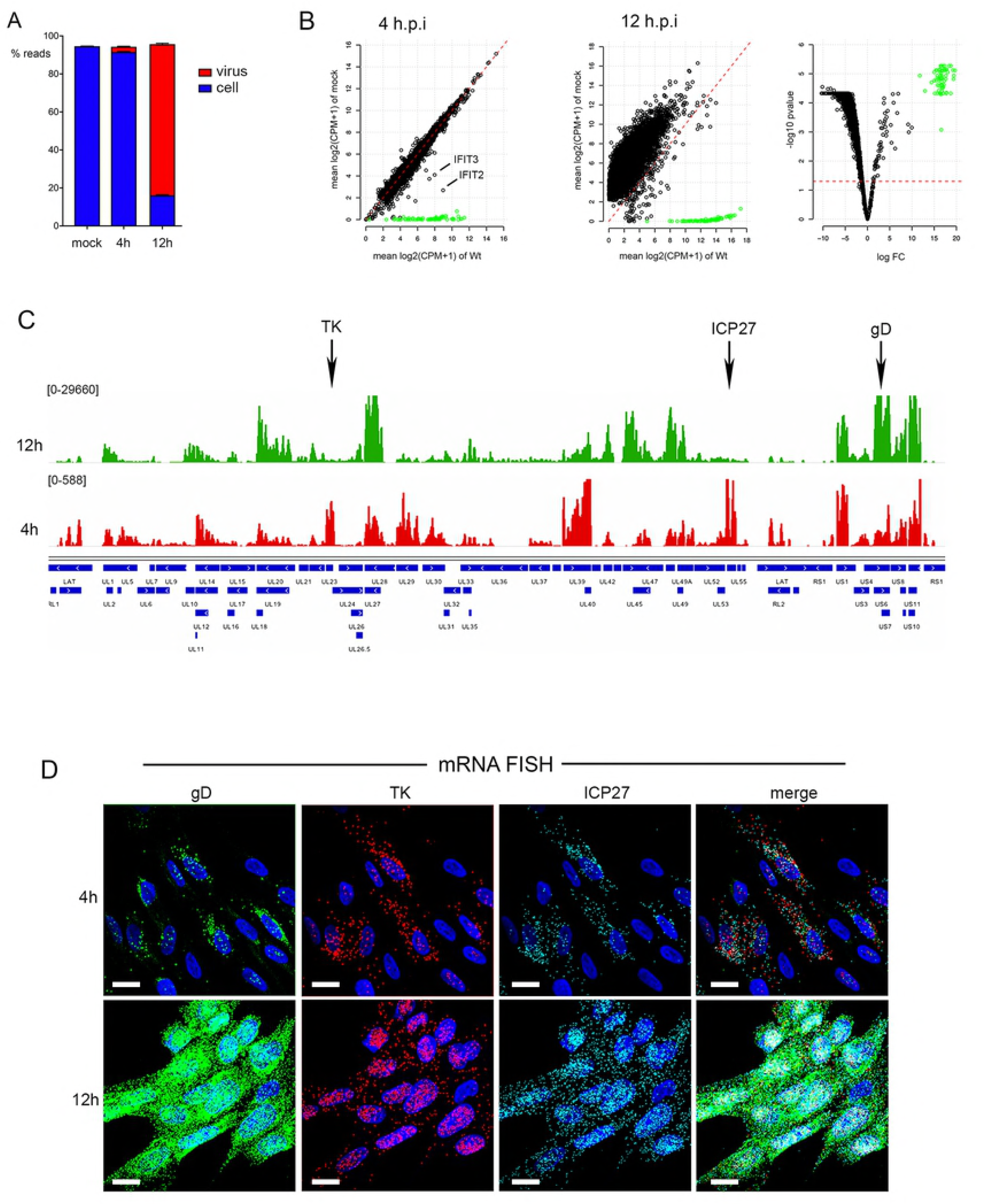
qRT-PCR of vhs-induced reduction in cellular transcript levels correlates with single cell mRNA FISH. (A) & (B) HFFF cells were left uninfected or infected with HSV1 Wt or Δvhs viruses at a multiplicity of 2. Total RNA was purified at 16 hours and subjected to qRT-PCR for transcripts identified as being susceptible to vhs activity (A) or for ISG transcripts (B). The mean and ± standard error of the data is given from one representative experiment (n=3). Statistical analysis was carried out using an unpaired, two-way student’s t test. **, p < 0.01. ***, p < 0.001. ****, p < 0.0001. (C) HFFF cells grown in chamber slides were left uninfected (mock) or infected with Wt or Δvhs viruses at a multiplicity of 2 and fixed at 16 hours. mRNA FISH was carried out for serpin E1, GLUL1 or IFIT1 (green). Nuclei were counterstained with DAPI (blue). Scale bar = 20 μm.

A potential explanation for variation in transcript susceptibility to vhs could be cell-to-cell variation in vhs activity or response. We monitored the transcripts for serpin E1 and GLUL - shown by RNAseq to be reduced at 12 hours in Wt infected cells by log_2_ FC of −7.45 and −0.9 respectively - by mRNA FISH of uninfected, Wt infected and Δvhs infected HFFF at 16 hours. The serpin E1 transcript signal was clearly decreased in Wt compared to mock or Δvhs infected cells (Fig 3C, serpin E1). By contrast, the GLUL transcript signal was maintained in infected cells compared to uninfected cells (Fig 3C, GLUL), providing further evidence for differential susceptibility of cell mRNAs to vhs activity. We also investigated IFIT1 mRNA levels by mRNA FISH confirming that the number of IFIT1 transcripts was increased in Δvhs infected cells compared to mock or 12-hour Wt infection (Fig 3C, IFIT1). Taken together, these data indicate that our RNAseq data correlates with results obtained by both qRT-PCR and mRNA-FISH.

### Translational shutoff in cells infected with HSV1 lacking the VP22 gene is not a consequence of overactive vhs

We next assessed the overactivity of vhs in Δ22 infections, by comparing the equivalent RNAseq libraries for Δ22 infected HFFF to the Wt infected libraries (S3 to S6 Tables), with differential expression plotted as scatter plots. Mapping the reads obtained for each library to the human and HSV-1 transcriptome indicated that the library composition of Δ22 infected cells was similar at 4 h but comprised a smaller percentage of virus transcripts at 12 h (Fig 2A). Unexpectedly, we found that there was little difference in the relative transcriptomes of Wt and Δ22 at either early or late times, with scatter plots of the total infected cell transcriptome showing limited difference in cellular (black circles) or virus (green circles) transcript levels (Fig 4B; S5 & S6 Tables. See also Fig S4). With our transcriptomic study selecting only two time points, we reasoned that we may have missed important differences between Wt and Δ22 infected cells, and hence a time-course of infection was carried out on Wt, Δ22 and Δvhs infected HFFF cells to determine changes in representative transcript levels over time. Relative changes of upregulated (IFITs 1 & 2), relatively insensitive (RPLP0 & GAPDH) and hypersensitive (MMP1 & MMP3) transcripts were then measured by qRT-PCR, with the Log_2_ FC plotted over time. In all three infections, the IFIT transcripts rose in abundance between 2 and 4 h, and in the absence of vhs, these antiviral transcripts were maintained at a high level throughout infection (Fig 4C, Δvhs). However, in Wt infection they began to decline almost immediately, while in Δ22 infection the levels kept rising and only started to drop around 8 h (Fig 4C, upregulated transcripts). Likewise, for vhs sensitive transcripts, degradation was also delayed in Δ22 compared to Wt infection, with degradation beginning at 8h and 4h respectively (Fig 4C). It was only the insensitive transcripts that appeared to be affected similarly in Wt and Δ22 infections (Fig 4C, low). The delayed activity of vhs in Δ22 infected cells was also confirmed by mRNA FISH of serpin E1 shown above (Fig 3C) to be greatly reduced during Wt infection, revealing that it was still present at uninfected cell levels 8h after infection with Δ22, but had been degraded by 16h, whereas the transcript was still abundant in Δvhs infection after 16 h (Fig 4D). These results suggest that up to 16h, vhs is not universally overactive against cellular transcripts in the absence of VP22.

**Fig 4.**
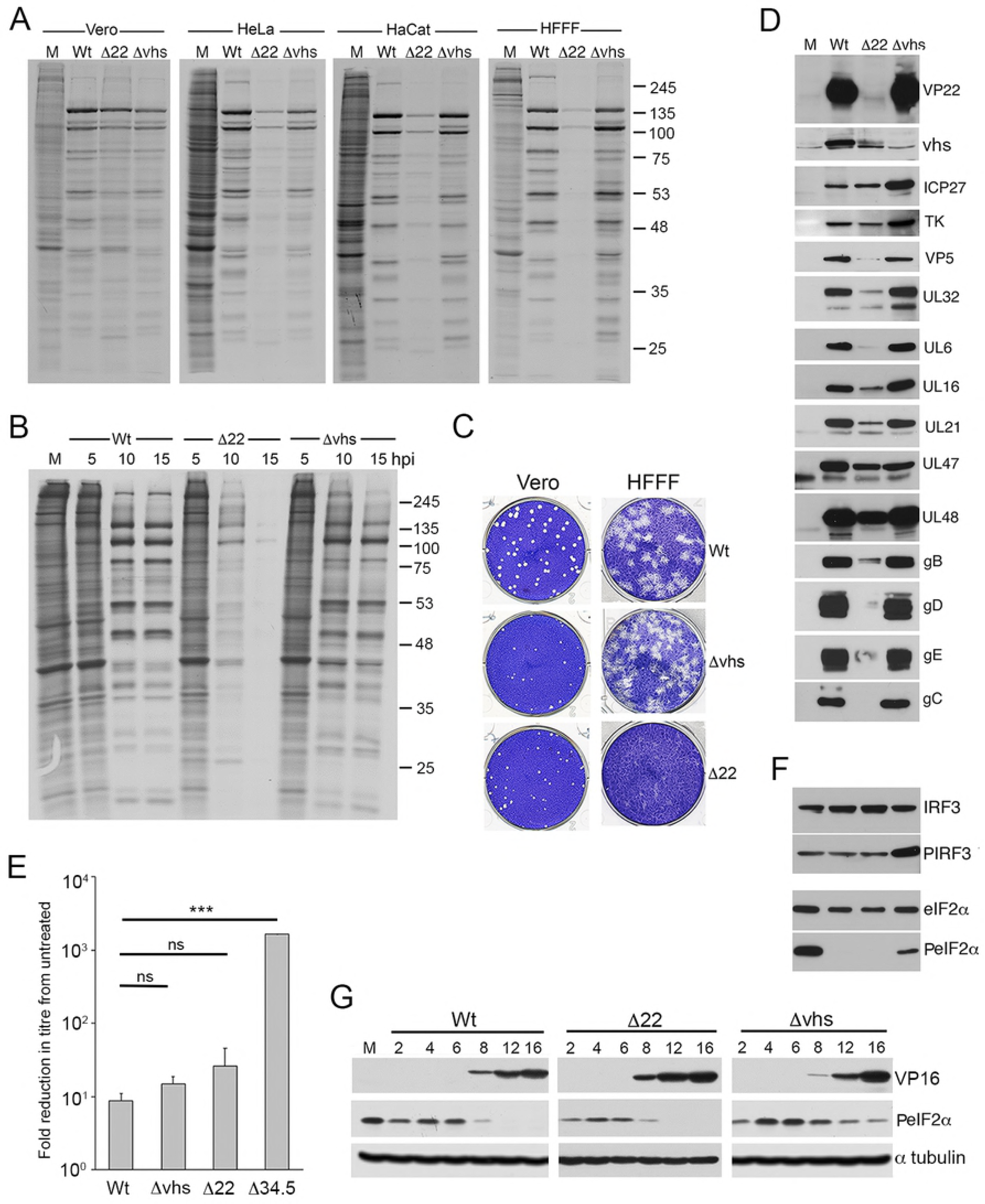
Vhs activity against cellular transcripts is delayed in Δ22 infected HFFF cells. (A) & (B) Dual transcriptomic analysis of HFFF cells infected with the Δ22 virus at a multiplicity of 2 was carried out alongside the analyses presented in Fig 2. (A) Proportion of reads mapped in each condition to either the human (blue) or HSV1 (red) transcriptome.(B) Differential expression analysis of cell and virus transcripts comparing Wt (y-axis) to Δ22 (x-axis) at 4 and 12 h after infection. Differences in the number of reads mapped to cell (black circles) and virus (green circles) transcripts were plotted as scatter plots. (C) HFFF cells were infected with Wt, Δ22 or Δvhs viruses at a multiplicity of 2, and total RNA was harvested at the indicated times (in hours). qRT-PCR was carried out for the indicated cell transcripts. Transcript levels are expressed as the log_2_ FC to mock (ΔΔCT) over time. The mean and ± standard error of the data is given from one representative experiment (n=3).(D) HFFF cells were infected with Wt, Δ22 or Δvhs viruses at a multiplicity of 2 were fixed at 8 or 16 hours after infection and processed for mRNA FISH with a probe specific for the cellular transcript for serpin E1. Nuclei were counterstained with DAPI. Scale bar = 20 μm. HFFF cells were infected with Wt HSV1 at a multiplicity of 2. (E) At 6 hours, the cells were either harvested for total RNA, or actinomycin D (5 μg/ml) was added and the infection left for a further 4 hours before harvesting total RNA. qRT-PCR was carried out on all samples for the indicated cell transcripts, with results expressed as the log_2_ FC to the sample harvested at 6 hours (ΔΔCT). The mean and ± standard error of the data is given from one representative experiment (n=3). Statistical analysis was carried out using an unpaired, two-way student’s t test. ns, p > 0.05. ***, p < 0.001. (F) HFFF cells were pre-treated in the absence or presence of cycloheximide (100 μg/ml) for 1 hour prior to infection with Wt HSV1 at a multiplicity of 2. Total RNA samples were purified at the indicate times and subjected to qRT-PCR for upregulated transcripts (IFIT1 and IFIT2), hypersensitive transcripts (MMP1 and MMP2) and virus transcripts (ICP27 and gC). For cell genes, the transcript levels are expressed as the log_2_ FC to mock, and for virus genes the are expressed as log_2_ FC to 2 hours. The mean and ± standard error of the data is given from one representative experiment (n=3). Statistical analysis was carried out using an unpaired, two-way student’s t test. ns, p > 0.05. *** p < 0.001.

In the experiments thus far described we had measured overall transcript levels rather than degradation rates alone. One possible explanation for the range of susceptibility to vhs could be that the variable level of loss in the cells reflects the balance between transcription and degradation of these transcripts. We therefore tested the relative loss of two vhs-insensitive transcripts (RPLP0 and GAPDH) and two vhs-sensitive transcripts (MMP1 and MMP3) in cells that had been incubated for 4 h with Actinomycin D to inhibit transcription from 6 h onwards. In uninfected cells, none of the transcripts tested were significantly altered in the presence of Act D, suggesting that they are all relatively stable over a 4h time period (Fig 4E, mock). The same experiment carried out in Wt infected cells indicated that even in the absence of ongoing transcription the variable susceptibility of the tested transcripts to vhs degradation held true, with the relative loss of highly susceptible transcripts being greater than those that are relatively resistant (Fig 4E, Wt). By contrast, the high susceptibility transcripts were not significantly degraded in Δ22 infected cells at this time (Fig 4E), a result that is consistent with the relative changes seen in the time course at this time of infection (Fig 4C) and correlates with the low level of vhs protein present in Δ22 infected cells (Fig 1D).

It is clear from the above data that the three groups of transcripts that we had identified – upregulated, insensitive and hypersensitive – behaved differently in their response to vhs activity in Wt infection. One potential explanation for such variation is that both incoming vhs from the virion and newly synthesized vhs could be differentially contributing to Mrna degradation over the first few hours of infection. To test the relative effect of incoming vhs, HFFF cells were infected with Wt HSV1 following pre-treatment with cycloheximide (CHX) to block protein translation, such that any mRNA degradation detected would be a consequence of vhs delivered by the virion. Total RNA samples were harvested at times up to 10 h and transcript levels for representative genes compared to those found in non-treated cells. qRT-PCR for the IE ICP27 transcript and the L gC transcript showed that, as expected, transcription of ICP27 occurred but was reduced while transcription of gC was blocked in the presence of CHX and hence the absence of ongoing infection (Fig 4F, right-hand panel). The hypersensitive MMP1 and MMP3 transcripts underwent slow degradation in CHX-treated cells, but as early as 2 hours after infection this was enhanced during active infection (Fig 4F, central panel). By contrast, the upregulated IFIT1 and IFIT2 transcripts which began to rise only by 4 hours were refractory to the activity of incoming vhs and required active infection to be degraded (Fig 4F, left-hand panel).

Bearing in mind that vhs can act against virus as well as cell transcripts, and that it is this effect that is predicted to cause translational shutoff that is detrimental to the virus, it is noteworthy that the relative levels of virus transcripts in the 12 h RNAseq data showed that there were only small differences in the virus transcriptome between Wt and Δ22 (Fig 4B, green circles). Nonetheless, those transcripts that were less abundant in Δ22 infection were predominantly those that encode L structural proteins (S6 Table & S5 Fig). To get a clearer picture of the effect of deleting VP22 on the levels of virus transcripts over time, we analysed the same time-course described in Fig 4 by qRT-PCR for a number of virus transcripts representing IE (ICP27 and ICP22); E (TK); and L (VP22, vhs and gC) genes. In this case, the ΔCT values were plotted to compare the relative expression of the individual transcripts in each virus at the denoted time points, and to determine how each mRNA changed over time in the presence and absence of VP22 and vhs. For all transcripts, there was a modest but consistent increase in the ΔCT value of all transcripts tested (ie a drop in transcript level) in the Δ22 infected cells compared to Wt at each time point (Fig 5A, compare Wt and Δ22) confirming the single point transcriptomic analysis. Between 8 and 16 h there was little difference in the ΔCT values of the L transcripts tested in Wt and Δvhs infected cells, but there was an increase for IE and E transcripts in Wt infection, with ICP27, ICP22 and particularly TK maintained at a high level at later times in the absence of vhs (Fig 5A). This agrees with studies from others [25] and likely reflects the fact that vhs degrades these mRNAs in Wt infections. Indeed, the drop in these transcript numbers in Wt infected cells correlated with the timing of vhs expression and activity from 6 h onwards. By contrast, in Δ22 infected cells all transcripts tested were found to be around 2-fold lower than in Wt infected cells throughout the course of the entire infection (Fig 5A, compare Δ22 and Wt), with the exception of gC which was up to10-fold lower by 16 hours. Western blotting of relevant virus proteins through the same time course confirmed the expected expression levels of representative proteins, including enhanced ICP27 levels in the Δvhs infection, and extremely low levels of L proteins in Δ22 infection (Fig 5B).

**Fig 5.**
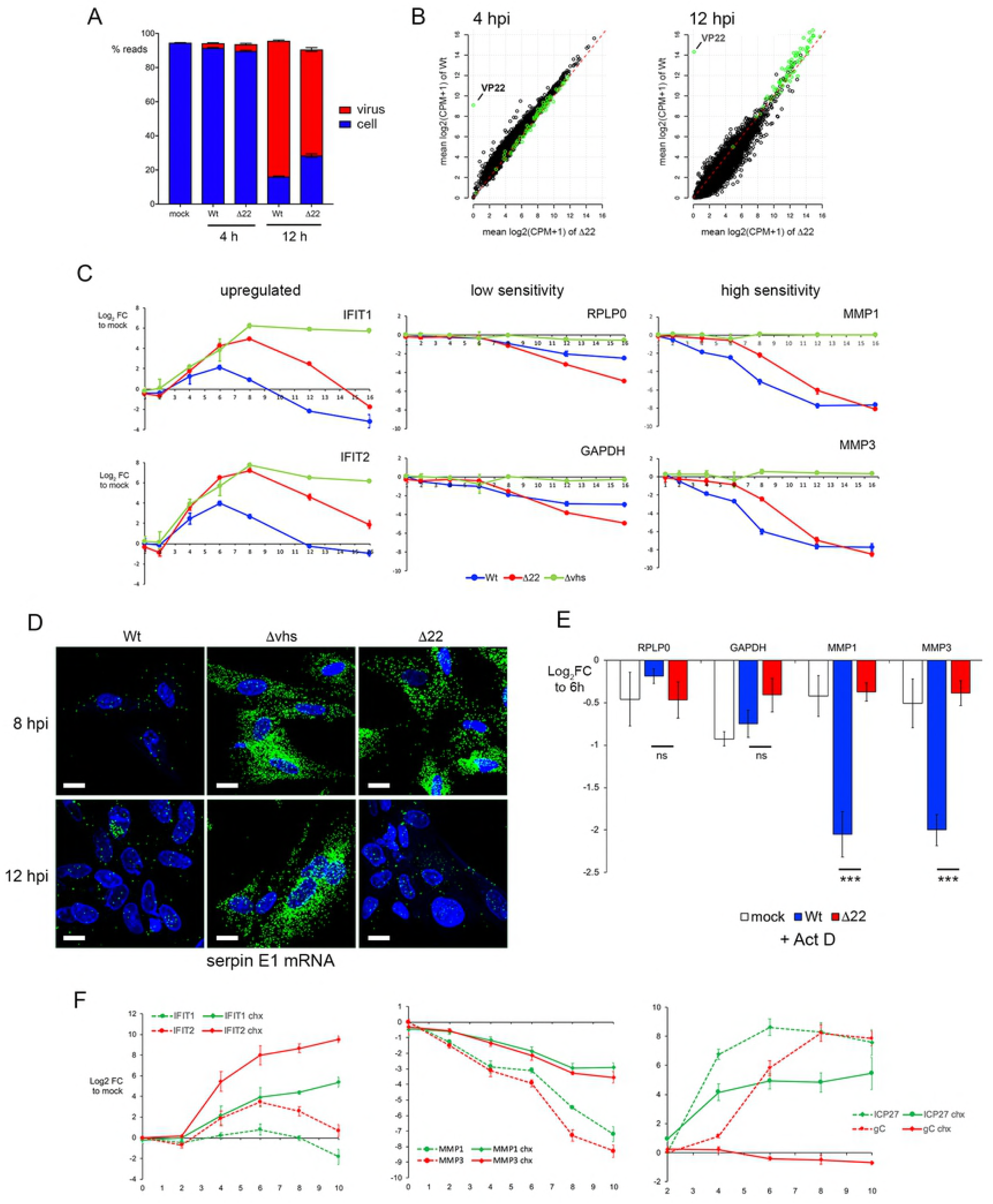
vhs exhibits differential activity against late virus transcripts in the absence of VP22. (A) The time course described in Fig 4C was analysed by qRT-PCR for representative IE (ICP27 and ICP22), E (TK) and late (VP22, vhs and gC) transcripts, with results represented as mean ΔCT values at each time point. (B) Total lysates harvested at the same time as the RNA samples in Fig 4C were analysed by SDS-PAGE and Western blotting with antibodies for the indicated proteins. (C) HFFF cells were infected with Wt HSV1 at a multiplicity of 2. At 6 hours, the cells were either harvested for total RNA, or actinomycin D (5 μg/ml) was added and the infection left for a further 4 hours before harvesting total RNA. qRT-PCR was carried out on all samples for the indicated virus transcripts, with results expressed as the log_2_ FC to the samples harvested at 6 hours (ΔΔCT). The mean and ± standard error of the data is given from one representative experiment (n=3). Statistical analysis was carried out using an unpaired, two-way student’s t test. ns, p > 0.05. * p < 0.05.** p < 0.01. *** p < 0.001.

The differences seen in virus transcript levels in the absence and presence of VP22 could be a consequence of differential transcription and/or degradation. Hence, we next tested the relative loss of a range of virus transcripts in cells that had been incubated for 4 h with Act D to inhibit transcription. The IE (ICP27 and ICP22) and E (TK) transcripts that were tested were all reduced around four-fold in these conditions, presumably reflecting their degradation by vhs, but there was no significant difference between the relative loss in Wt and Δ22 infected cells (Fig 5C). By contrast, while all L transcripts tested were equally susceptible to vhs as the earlier classes of transcript in the absence of VP22, these L transcripts were minimally reduced in Wt infection (Fig 5C). This indicates that VP22 differentially “protects” L transcripts from degradation by vhs.

### Translational shut-off in cells infected with HSV1 lacking the VP22 gene correlates with vhs-induced nuclear retention of L virus transcripts

Taken together with our results above demonstrating that vhs degradation activity is not vastly overactive in Δ22 infected cells, we reasoned that the extreme translational shutoff seen in the absence of VP22 may be a consequence of mRNA localisation rather than levels. To understand the behaviour of different classes of virus transcripts, we next examined the localisation of representative virus transcripts up to 12h in Wt infected cells – the IE ICP27 and ICP0 transcripts, the E TK transcript and the L vhs and gC transcripts. ICP27 and ICP0 were readily detectable at 2h and increased in numbers up to 8h. While ICP0 transcripts became obviously retained in the nucleus by 6h, an indication of the fact it is a spliced transcript and, as shown by others its nuclear export would be inhibited by the activity of ICP27 protein after DNA replication [4, 56], ICP27 transcripts were cytoplasmic up to 12h when some level of nuclear retention became obvious (Fig 6, ICP27 & ICP0). In the case of TK, a few transcripts were present in the nucleus at 2 hours indicative of transcription just starting, but these became more obvious in the cytoplasm by 4 hours and like ICP27 remained cytoplasmic until 12 h when a significant number of transcripts were present in the nucleus. By contrast, the L gC transcript was only detectable at 4 hours and remained entirely cytoplasmic up to 12 hours, despite the large number of transcripts present (Fig 6, gC). Interestingly for vhs, despite being classed as a L gene, small numbers of transcripts were detected in the nucleus as early as 2 hours and persisted until 8 hours when its numbers increased. However, even by 12 hours the vhs mRNA was predominantly nuclear (Fig 6, vhs).

**Fig 6.**
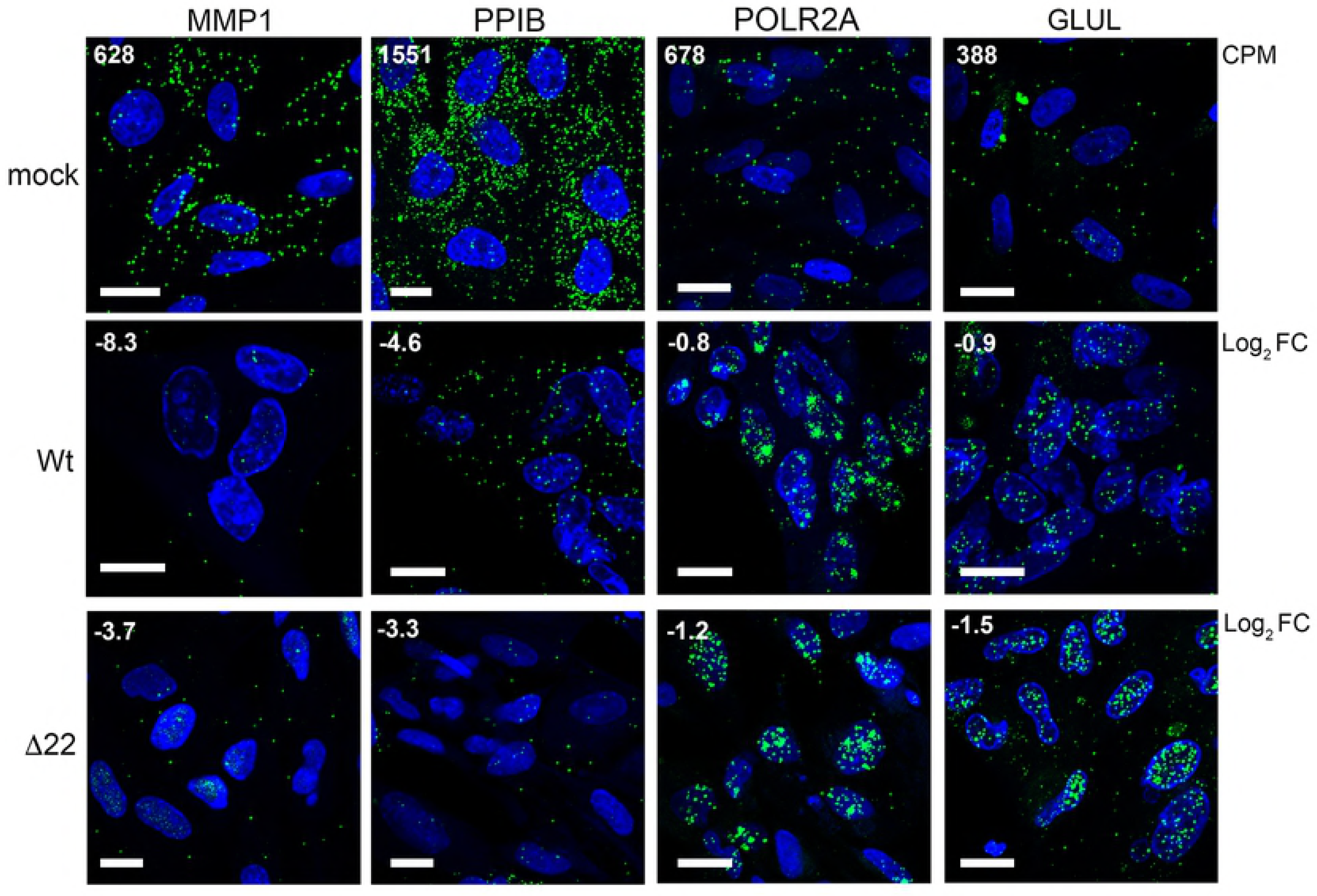
HSV1 transcripts exhibit differential subcellular localisation. HFFF cells grown in slide chambers were infected with Wt virus at a multiplicity of 2, fixed at 2, 4, 6, 8 or 12 hours after infection, and processed for mRNA FISH with probes to the IE transcripts, ICP27 and ICP0, the E transcript TK and the late transcripts vhs and gC (all in red). Nuclei were counterstained with DAPI (blue). Scale bar = 20 μm.

In light of the above results, it is notable that we have recently reported that when expressed in isolation, vhs causes the nuclear retention of its own transcript through a negative feedback loop [42]. Moreover, we also demonstrated the nuclear retention of co-expressed transcripts which correlated with nuclear localisation of the polyA binding protein (PABP) in transfected cells, in agreement with other studies [42, 43, 57, 58]. Here we found that Wt HSV1 infection of HFFF also altered the localisation of PABP from being exclusively cytoplasmic to being predominantly nuclear (Fig 7A). In Δvhs infected cells, PABP localisation remained cytoplasmic, confirming the role of vhs in its re-localisation to the nucleus (Fig 7A). Interestingly, the re-localisation of PABP to the nucleus was more pronounced in Δ22 infected cells where it was exclusively nuclear compared to Wt infection (Fig 7A), suggesting that this activity of vhs is greatly enhanced in HFFF cells in the absence of VP22. This led us to investigate the relative localisation of IE (ICP27), E (TK) and L (gC) transcripts by mRNA FISH at 8 and 16 hours after infection – later than the previous experiment but in line with our metabolic labelling studies - in Wt, Δvhs and Δ22 infected cells. At 8h, all three mRNAs exhibited similar cytoplasmic localisation regardless of the presence or absence of vhs or VP22 (Fig 7B, 8 hpi). By contrast, there was a striking difference at 16 hpi, when in Wt infected cells, ICP27 and TK mRNAs were substantially localised to the nucleus as implied by the data presented in Fig 6, while gC was predominantly cytoplasmic. However, in Δvhs infected cells, all 3 transcripts were entirely cytoplasmic, suggesting that vhs activity is required for the differential localisation of virus transcripts seen in Wt infected cells. This provides evidence that vhs activity not only degrades IE and E transcripts but causes their nuclear retention, a mechanism that would by definition efficiently block translation of these mRNAs at late times when they are not required. Moreover, L transcripts appear to be spared this effect of vhs thereby allowing them to be efficiently translated at late times. Intriguingly, not only were the IE and E transcripts predominantly nuclear by 16h in Δ22 infected cells, more so than in Wt infected cells, but also the L gC transcript was almost entirely retained in the nucleus (Fig 7B, 16 hpi), a situation that also held true for the L VP16 transcript (Fig 7C). In addition, mRNA FISH of the spliced IE transcript showed that it was retained in the nucleus of all three infections including the Δvhs infection, albeit more so in the absence of VP22 (Fig 7C). As such, our results demonstrate that in HFFF cells, VP22 is required for the cytoplasmic localisation and hence translation of L virus transcripts in the presence of active vhs.

**Fig 7.**
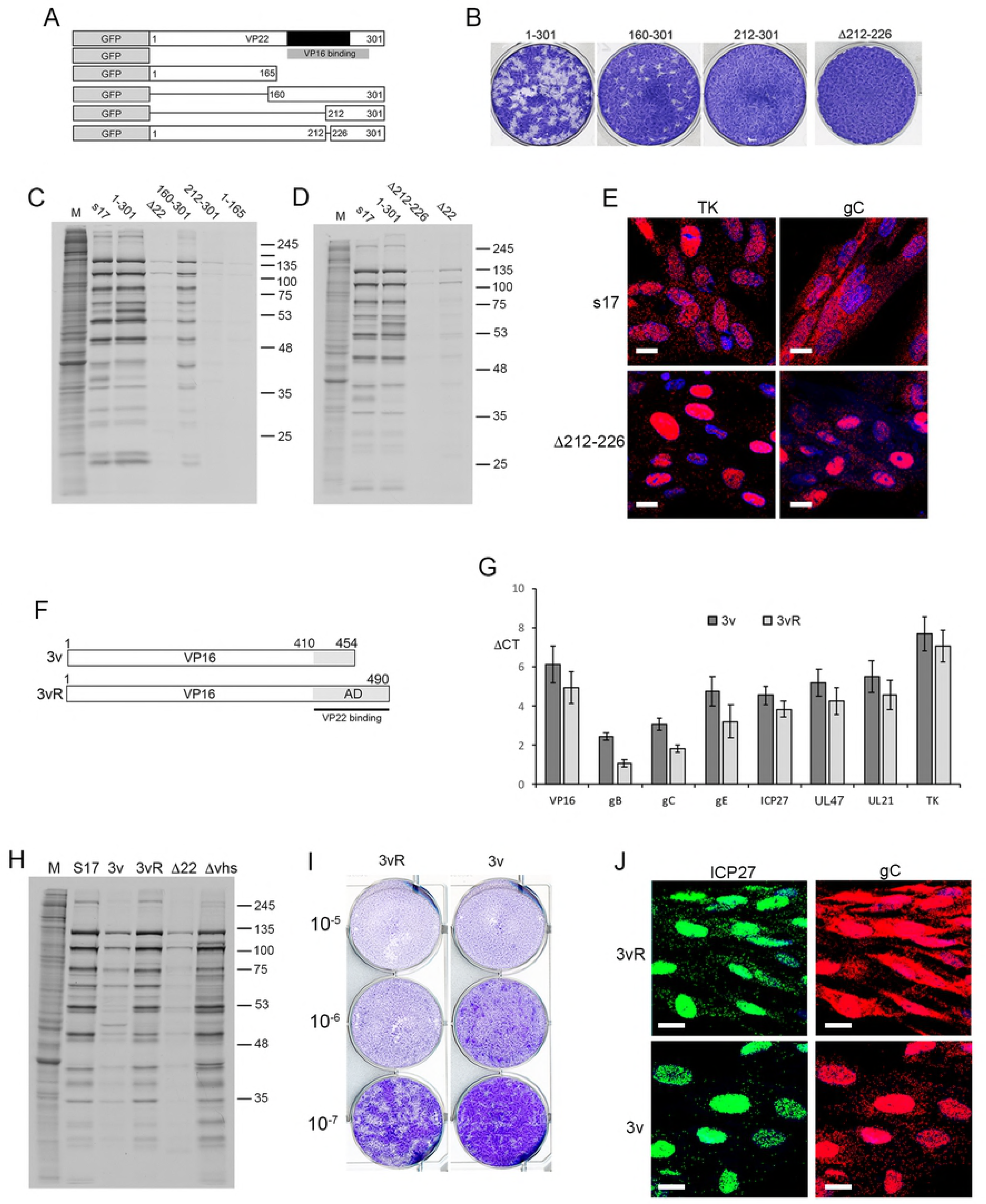
Nuclear retention of late virus transcripts in the absence of VP22. (A) HFFF cells infected with Wt, Δvhs or Δ22 viruses at a multiplicity of 2 were fixed 12 hours after infection and processed for immunofluorescence with an antibody to PABP (green). Nuclei were counterstained with DAPI (blue). (B) HFFF cells infected with Wt, Δvhs or Δ22 viruses at a multiplicity of 2 were fixed at 8 or 16 hours after infection and processed for mRNA FISH with probes specific to an IE (ICP27), E (TK) or late transcript (gC) (all in red). Nuclei were counterstained with DAPI (blue). (C) As for B, but cells fixed at 16 hours were processed for mRNA FISH with a probe specific to the late transcript VP16, or the spliced IE transcript ICP0 (both in red). Nuclei were counterstained with DAPI (blue). Scale bar = 20 μm.

### Cell transcripts insensitive to vhs degradation exhibit differential nuclear retention in the presence or absence of VP22

In the absence of VP22 we have shown that all virus transcripts tested thus far are retained in the nucleus. Given that this retention correlates with a limited but consistent reduction in transcript levels, we hypothesized that the relative level of reduction in cellular transcripts may reflect the degree of nuclear retention exhibited by those transcripts. Using mRNA FISH, we examined the localisation of two transcripts hypersensitive to vhs (MMP1 & PPIB) and two relatively insensitive transcripts (POLR2A & GLUL) in Wt and Δ22 infected cells at 12 h in comparison to uninfected cells. Importantly, the numbers of each transcript detected in uninfected cells correlated well with the average CPM found in our uninfected RNAseq data (Fig 8, mock, CPM number in top left-hand corner). Moreover, the level of depletion detected in both Wt and Δ22 infected cells also correlated with our RNAseq data (Fig 8, Wt & Δ22, Log_2_ FC shown in top left-hand corner), with MMP1 and PPIB transcripts reduced greatly in numbers, but little difference determined in POLR2A & GLUL. Strikingly, both POLR2A and GLUL transcripts were retained in the nucleus of Wt infected cells, a feature that was amplified in Δ22 infected cells (Fig 8), suggesting that the vhs-dependent block to nuclear export of virus transcripts also resulted in the nuclear retention of these cellular transcripts. Such nuclear retention would by definition minimise the susceptibility of those nuclear-entrapped transcripts to further degradation by vhs resulting in only a minimal loss of transcript. By contrast, those transcripts that were highly degraded were not retained in the nucleus, reflecting their degradation prior to nuclear retention activity.

**Fig 8.**
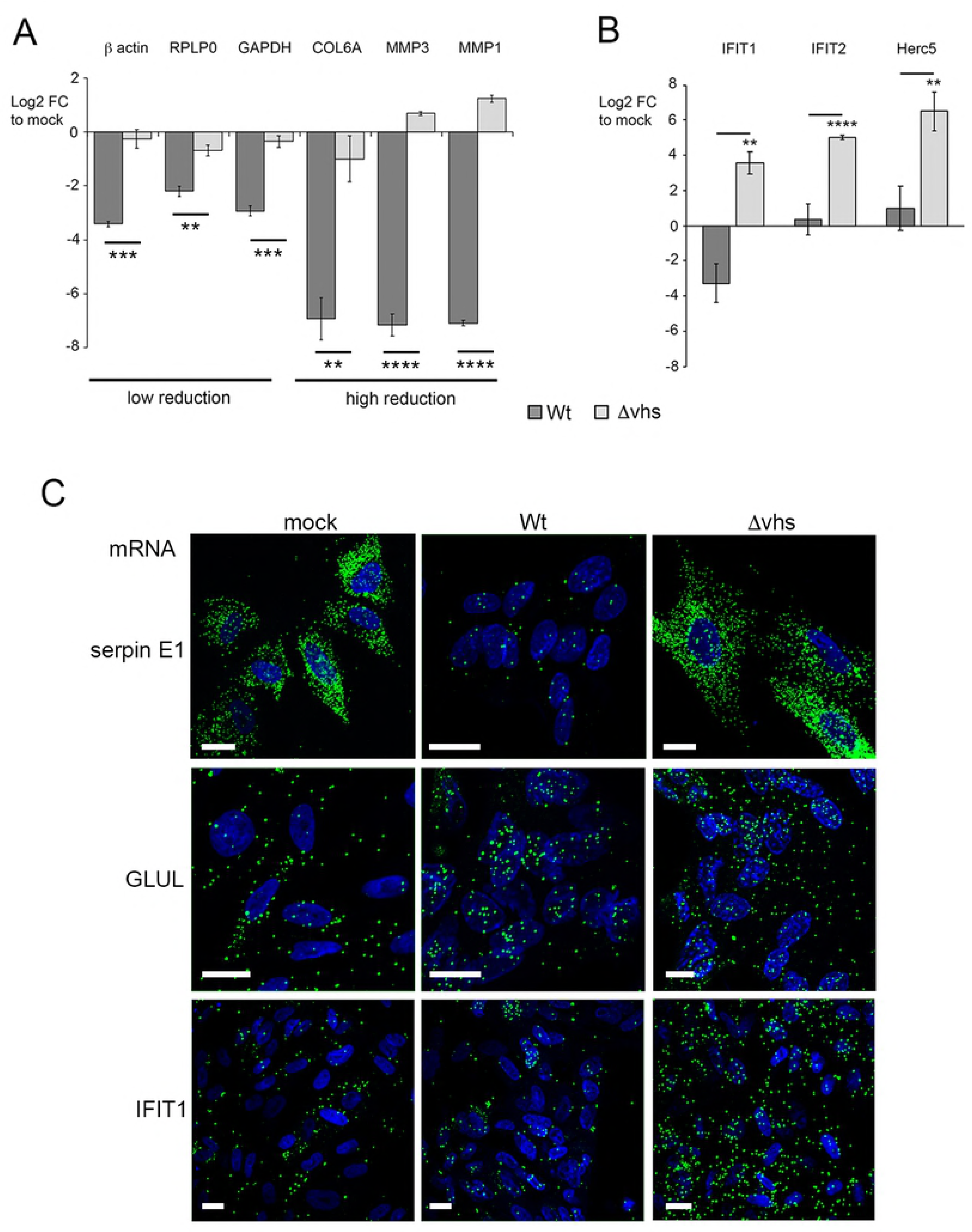
Cell transcripts insensitive to vhs activity are retained in the nucleus of infected cells. HFFF cells infected with Wt or Δ22 viruses at a multiplicity of 2 were fixed at 12 hours after infection and processed for mRNA FISH with probes specific to two vhs hypersensitive cell transcripts, MMP1 and PPIB, and two vhs-insensitive cell transcripts, POLR2A and GLUL (all in green). Nuclei were counterstained with DAPI (blue). Scale bar = 20 μm. CPM = average read counts per million in uninfected RNAseq libraries. Log_2_ FC = log_2_ fold change in Wt and Δ22 infected RNAseq libraries at 12 hours as determined by bioinformatic analysis.

### Export of the infected cell transcriptome from the nucleus is enhanced by VP22 binding to VP16

Given that our results here suggest that the outcome of virus infection in the absence of VP22 is the nuclear retention of L transcripts and translational shutoff of L proteins, we next tested the requirement for the conserved domain of VP22 – ther region that contains the VP16 binding domain - in this phenotype using a panel of previously described recombinant viruses expressing deletion mutants of VP22 fused to GFP [59, 60] (Fig 9A). Plaque assays on HFFF cells revealed that insertion of GFP at the N-terminus of VP22 had little effect on plaque formation, while the C-terminal half of the protein containing the conserved domain of VP22 (160-301) was sufficient to maintain plaque formation. However, further deletion into this region (212-301) or deletion of only 12 residues within this domain (Δ212-226) prohibited plaque formation in these cells (Fig 9B). Likewise, metabolic labelling indicated that the same mutant viruses that failed to form plaques on HFFF resulted in translational shutoff compared to GFP-22 expressing virus or virus expressing the C-terminal half of VP22 (Fig 9C & D). Furthermore, mRNA FISH of cells infected with the Δ212-226 virus and fixed at 16 hours revealed that this small deletion in VP22 was sufficient to cause nuclear retention of gC transcripts thereby producing a phenotype equivalent to deletion of the entire VP22 protein (Fig 9E).

**Fig 9.**
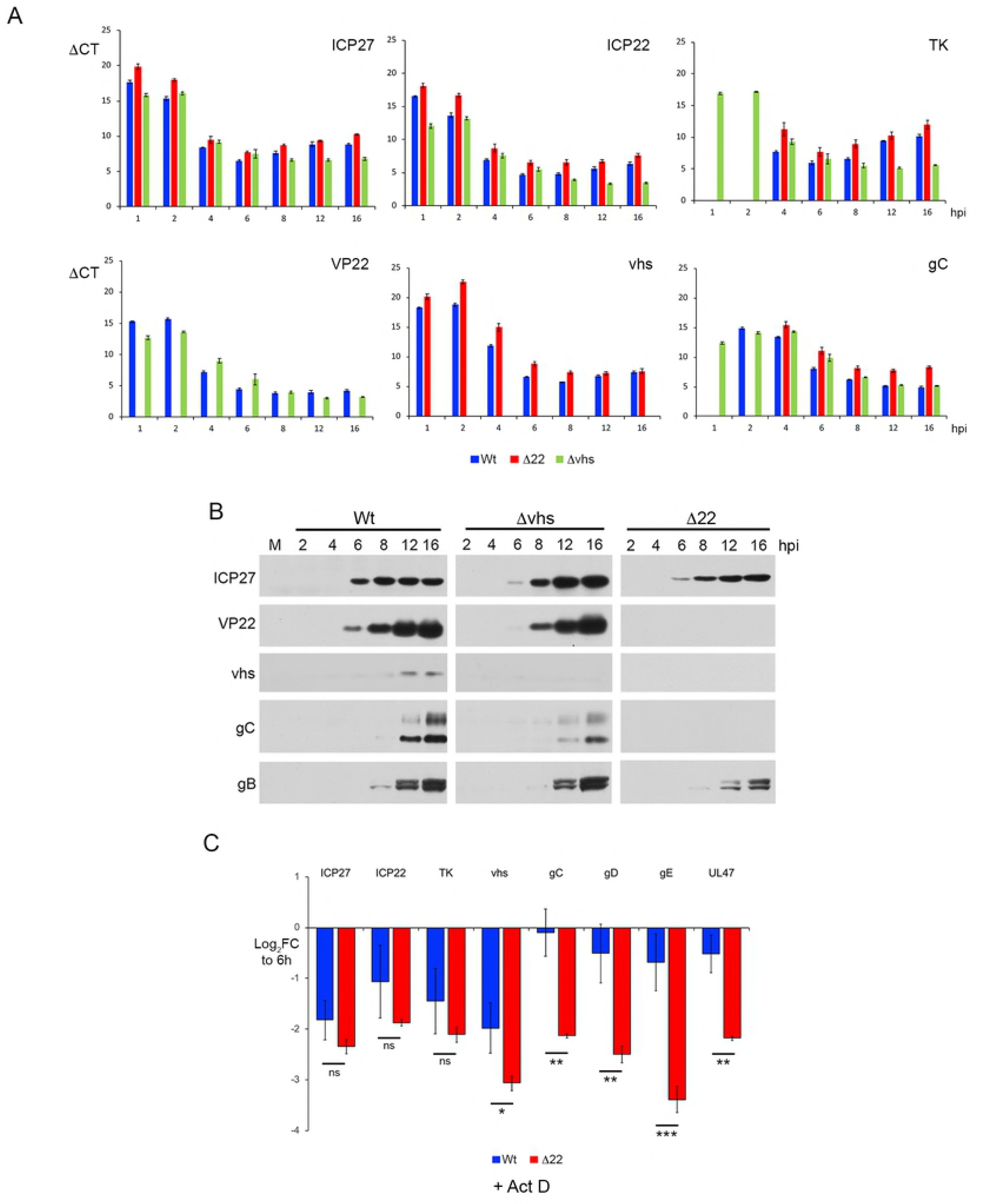
Cytoplasmic localisation of late transcripts is enhanced by VP22 binding to VP16. (A) Line drawing of VP22 variants expressed as GFP fusion proteins in virus infection. Black box indicated the conserved domain of VP22, grey box indicates the region required for VP16 binding. (B) Plaque formation of viruses shown in A on HFFF cells. (C) & (D) HFFF cells were infected with Wt (s17), Δ22 or the viruses shown in A at a multiplicity of 2, were metabolically labelled with [35S]-methionine 15 hours after infection. Cells were lysed and analysed by SDS-PAGE and autoradiography. (E) HFFF cells infected with Wt (s17) or Δ212-226 at a multiplicity of 2 were fixed at 16 hours and subjected to mRNA FISH with probes specific for the IE transcript ICP27 or the late transcript gC (both in red). Nuclei were counterstained with DAPI (blue). Scale bar = 20 μm. (F) Line drawing of the variant of VP16 (Δ454-490) expressed in the 3v virus based on the KOS strain, together with its rescue virus 3vR. The grey box indicates the C-terminal activation domain (AD) of VP16, the black line indicates the region of VP16 required to bind VP22. (G) HFFF cells infected with the viruses shown in F at a multiplicity of 2 were harvested for total RNA at 16 hours and analysed by qRT-PCR for the indicated transcripts. Results are represented as ΔCT values. (H) HFFF cells infected with Wt (s17), Δ22, Δvhs or the viruses shown in F at a multiplicity of 2, were metabolically labelled with [35S]-methionine 15 hours after infection. Cells were lysed and analysed by SDS-PAGE and autoradiography. (I) The 3v and 3vR viruses were titrated onto HFFF cells and plaques fixed and stained with crystal violet 5 days later. (J) HFFF cells infected with 3v or 3vR viruses at a multiplicity of 2 were fixed at 16 hours and subjected to multiplex mRNA FISH with probes specific for the IE transcript ICP27 (green) and the late transcript gC (red). Nuclei were counterstained with DAPI (blue). Scale bar = 20 μm.

VP22 and vhs both bind to VP16 to form a trimeric complex [31, 32, 42] through which they are jointly proposed to quench vhs activity [36, 61]. The C-terminal half of VP22 shown above to be required to rescue vhs-induced nuclear retention of L transcripts has been reported to be involved in many activities including its binding to VP16, and hence the only role of this region may be to bring VP22 into the VP16-vhs complex. In an attempt to discriminate between a requirement for this region in direct VP22 activity rather than simply binding to VP16, we utilised a previously described virus that lacks the C-terminal 36 residues of VP16 that are required to bind to VP22 [42, 62] as outlined in Fig 9F. Comparison of representative transcript levels by qRT-PCR between this mutant virus (3v) and its revertant (3vR), indicated that similar to a Δ22 infection, all transcripts tested were present at around 2-fold lower levels in the absence of the VP16-VP22 complex (Fig 9G). Nonetheless, metabolic labelling studies (Fig 9H), plaque assays on HFFF (Fig 9I) and mRNA FISH of the gC transcript (Fig 9J) indicated that the phenotype of the 3v virus was less extreme than the Δ22 virus in all assays, suggesting that while the interaction of VP22 with VP16 enhances the role of VP22, it is not essential. In summary, our data suggests that vhs, in combination with VP22 and VP16, co-ordinates the temporal expression of virus genes by retaining IE, E and cell transcripts in the nucleus while allowing the export of L transcripts, thereby ensuring that the translation of structural proteins is dominant for virus assembly at this time.

## Discussion

Many viruses encode endoribonucleases which promote the degradation of host mRNAs to block host gene expression, resulting in translational shutoff during infection [63]. However, the link between mRNA transcript degradation and translational shutoff has proved to be more complex than originally believed, primarily because such endoribonucleases have the potential to act on virus as well as cell transcripts. In this study we have used a combination of transcriptomics and single cell mRNA analysis to not only characterise in detail the activity of the HSV1 vhs endoribonuclease, but to determine the role of one of its cofactors, VP22, in regulating vhs behaviour. RNAseq studies were performed at 4 and 12 hours after infection of HFFF cells enabling us to determine the fate of some 11,000 cellular transcripts and all virus transcripts at early and late times of infection. Two previously reported microarray studies covered a similar time-frame of Wt and Δvhs virus infection but both focused only on transcripts that were upregulated during infection, and in particular those that were activated as part of the host innate immune response [13, 29]. By contrast, two more recent RNAseq based transcriptomic studies looked at the overall changes in ∼11,000 cellular transcripts [64, 65], but only went as far as 8 hours after infection, and therefore missed the greatest effect of vhs activity which as we have shown here occurs between 8 and 12 hours. Of note, because of the vast change in the relative content of the cell and virus components of the total transcriptome by 12 hours, and hence changes in the absolute amount of each component of the transcriptome, we were careful to normalise our data using ERCC control transcripts to spike the RNA samples prior to library production [66, 67]. Subsequent validation of a range of transcripts by qRT-PCR provided confidence in the RNAseq differential expression analyses of cellular and virus genes under different conditions.

*In vivo*, the main role of vhs on cellular transcripts is believed to be the degradation of interferon-induced transcripts that express antiviral proteins [13-17, 19]. Human fibroblasts were chosen for the study here due to the extreme translational shutoff phenotype of our Δ22 virus in these cells compared to Vero cells, pointing to cell-type variation in the requirement for VP22. Given that HFFF are primary human cells that can elicit a full antiviral interferon response pathway, whereas Vero cells are unable to express interferon [48] we reasoned initially that the differential phenotype in the absence of VP22 may be due to antiviral responses expressed in HFFF but not Vero cells. However, our results indicated that HFFF cells respond similarly to Δ22 infection compared to Wt infection, while mounting a strong innate immune response to Δvhs infection as expected [13]. Our studies show that the majority of the ∼100 upregulated genes in infected HFFF cells at 4 hours were classified as ISG transcripts, with at least two of these (IFIT1 and IFIT2) shown by our more detailed studies to be activated between two and four hours after infection. During infection, these induced transcripts declined over the course of roughly 12 hours in a vhs-dependent fashion, while IRF3 phosphorylation was enhanced in the absence of vhs, reflecting the signalling events that occur when vhs is not present. Intriguingly our studies with cycloheximide indicated that while it was components of the incoming virus that activated the innate immune response to upregulate ISG expression, only newly synthesized vhs protein and not virion delivered vhs was able to degrade thse activated ISG transcripts. In the absence of VP22, this ISG degradation was delayed but nonetheless occurred, a situation that mirrors vhs activity on other susceptible cellular transcripts, and may be a consequence of the low level of vhs expressed in Δ22 infected cells [37, 42]. Thus, our results do not concur with a recently published study that suggests that VP22 itself inhibits cellular DNA sensing through cGAS/STING, and that HSV1 infection in the absence of VP22 fails to inhibit interferon production [68].

Two major features - that are likely to be linked - have emerged from our studies on vhs. First, our RNAseq data revealed the differential degradation of cell transcripts ranging from those that were reduced by log_2_ FC of 10 (over 1000-fold), to those that were hardly altered. Such a diverse effect of vhs was not a consequence of transcript abundance or, at least for the transcripts that we tested, transcription dynamics. Detailed work from the Roizman lab has shown previously that vhs selectively spares a small number of cellular transcripts from degradation, a feature they have suggested could be sequence-specific [69, 70]. However, the broader picture as presented here encompasses the effect of vhs on 11,000 cellular transcripts and suggests there is a continuum of susceptibility to vhs activity. A possible explanation for differential effects of vhs on the cellular transcriptome could be that vhs preferentially targets transcripts in particular cellular locations or subsets of ribosomes. While the detail of the transcripts that are resistant or susceptible to vhs will not be considered further here, it is interesting to note that the majority of those that were efficiently depleted (> Log_2_ FC of −5) encode membrane and secreted proteins, suggesting that vhs may have enhanced activity for transcripts translated on the endoplasmic reticulum.

Nonetheless, we suggest that the differential effects of vhs are likely to be linked to the second major feature of vhs activity, its effect on the relative nuclear-cytoplasmic ratio of the infected cell transcriptome. We have shown for the first time that vhs expression from around 8 hours onwards of HSV1 infection causes the retention of multiple transcripts – cellular and viral - in the nucleus of the infected cell, thereby producing an effective mechanism of translational shutoff that does not require complete mRNA degradation. This single feature reveals the mechanism by which the virus efficiently regulates the transition from IE/E to L gene expression, as by selectively retaining IE/E but not L transcripts in the nucleus, translation is effectively switched from early to late phases. These results complement our previous study on vhs expressed by transient transfection [42], and add to a growing consensus that the activity of viral endoribonucleases in the cytoplasm affects events in the nucleus. In particular, work on the KSHV sox protein has shown that sox expression blocks RNA polymerase II activity on the cellular genome [71] and causes mRNA transcripts to become hyper-adenylated in the nucleus [43]. In KSHV and HSV1 infected cells, PABP – a protein that has a steady-state cytoplasmic localisation but shuttles between the cytoplasm and nucleus to bind polyadenylated mRNAs ready for export [72] – is released from mRNAs that have been degraded and is imported in to and accumulates in the nucleus in an endoribonuclease dependent fashion [42, 73, 74]. Here we found that in HFFF cells, PABP relocalised to the nucleus between 8 and 12 hours after infection correlating with the major expression of vhs. PABP was also more efficiently retained in the nucleus of Δ22 compared to Wt infected cells, which correlated with the fact that in the absence of VP22, every transcript tested underwent more efficient nuclear retention at this time. Such a phenotype suggests that the nuclear retention activity of vhs is non-selective and explains the profound translational shutoff seen in Δ22 infected cells even though the virus transcriptome was only minimally altered compared to Wt. Moreover, it reveals the true role of VP22 in the regulation of vhs activity, in that it rescues the cytoplasmic localisation of L transcripts in particular, rather than their hyper-degradation.

Our data indicates three phases to vhs expression and activity. First, vhs brought in by the virion begins to degrade highly susceptible cellular transcripts but does not act on stimulated ISG transcripts. Second, a low number of vhs transcripts detectable as early as 2 hours (as detected by mRNA FISH), and maintained at around 5 to 10 transcripts per cell up to 8 hours was able to express enough vhs protein to continue the degradation of highly susceptible transcripts and innate antiviral transcripts over and above the activity of incoming vhs protein. Of note, unlike other virus transcripts, these vhs transcripts were retained in the infected cell nuclei from the outset, in a manner similar to that detected in our studies of cells transiently expressing vhs [42]. Third, the transcription (and translation) of vhs was enhanced later in infection (∼ 8 hours) after which it caused the rapid nuclear retention of virus and insensitive cellular transcripts together with PABP, results that we had also previously observed in HeLa cells overexpressing vhs by transient transfection [42]. It is this third phase of vhs activity from which VP22 differentially protects L transcripts. These three waves of activity may indicate that a threshold of vhs protein and/or activity needs to be reached before nuclear retention occurs.

An obvious question is therefore how our results fit with the well-characterised role of ICP27 in late gene expression, where it has been shown that ICP27 is required for cytoplasmic localisation of late transcripts by binding to them and facilitating their nuclear export [5-7]. Moreover, ICP27 has been implicated in the translation of L proteins [75-77] while vhs and ICP27 have been reported to interact on translating viral mRNAs [78]. Of note, VP22 has been characterised as an RNA-binding protein [79, 80] via its C-terminal domain shown here to be important for its activity on L-transcripts, and small amounts of it can be detected in the nucleus of infected cells [81], and it is therefore tempting to speculate that VP22 is somehow involved in ICP27-directed export of L transcripts. The fact that VP22 and vhs both interact with different regions of VP16 [31, 32] and the incomplete phenotype of a virus expressing a variant of VP16 that was unable to interact with VP22 (while still binding vhs) also suggest that VP22 may be brought into proximity with vhs via VP16, an event that could occur as vhs is brought to the mRNA during translation initiation on the ribosome. We therefore suggest that the biogenesis of RNA in HSV1 infected cells is co-ordinated by a combination of multiple virus factors that vary in a temporal fashion, and which together ensure that both viral and cellular mRNAs are in the right place at the right time to ensure the productivity of the infected cell is dedicated to make new virus particles. In summary, our results present a new outlook on the complex subject of herpesvirus gene expression, providing scope to understand and tease apart the relative contributions made by each of these proteins to RNA biogenesis, localisation and translation.

## Methods

### Cells and Viruses

HFFF, HeLa (both obtained from European Collection of Authenticated Cell Cultures - ECACC) and HaCaT (obtained from Prof J. Breuer) cells were cultured in DMEM supplemented with 10% foetal bovine serum (Invitrogen). Vero cells (obtained from ECACC) were grown in DMEM supplemented with 10% newborn calf serum (Invitrogen). Viruses were routinely propagated in Vero cells, with titrations carried out in DMEM supplemented with 2% human serum. HSV1 strain 17 (s17) was used routinely. The s17 derived VP22 deletion mutant (Δ22) and the vhs knockout virus (Δvhs) have been described before [28, 38]. HSV1 strain Kos with a deletion of the C-terminal 36 residues of VP16 (RP3v) and its revertant (RP3vR) have been described elsewhere [62] and were kindly provided by Steve Triezenberg (Van Andel Institute). The s17 derived ΔUL13 and ΔICP34.5 knockout viruses have been described previously [82, 83]. The construction of viruses expressing GFP- tagged VP22 (GFP1-301), and GFP-tagged VP22 subdomains (GFP192-301, GFP108-301, GFP1-212, GFP1-165 and GFPΔ213-226) has also been described before [59, 60, 84].

### Antibodies & reagents

VP22 (AGV031) and UL47 (5283) antibodies have been described elsewhere [80, 85]. Other antibodies used in this study were kindly provided by the following individuals: gD (LP14), VP16 (LP1) and gB (R69), Tony Minson (University of Cambridge); vhs, Duncan Wilson (Albert Einstein College of Medicine); gE (3114), David Johnson (Oregon Health and Science University, Portland, OR USA); UL16 and UL21 John Wills (Penn State University); TK, UL6 and UL32, Frazer Rixon, Centre for Virus Research, Glasgow. Other antibodies were purchased commercially - α-tubulin (Sigma), VP5 (Virusys), gC, phospho-eIF2α, eIF2α, IRF3, phospho-IRF3 (AbCam), PABP and ICP0 (Santa Cruz). Horseradish peroxidase-conjugated secondary antibodies were from Bio-Rad Laboratories. Actinomycin D (Sigma) was used at a concentration of 5 μg/ml. Cycloheximide (Sigma) was used at a concentration of 100 μg/ml.

### Plaque reduction assay

Vero cell monolayers were pre-treated with 1000 units/ml recombinant human IFN-ß (R&D Systems) for 24 hours prior to infection with serial dilutions of wild-type HSV-1 (s17) or relevant mutants derived from this strain. The titres were determined by counting the number of plaques after 96 hours in the presence of human serum and results were expressed as a ratio of the titres observed in the presence or absence of interferon.

### Metabolic labelling of infected cells

Cells grown in 3cm dishes were infected at a multiplicity of 2, and at indicated times were washed and incubated for 30 mins in methionine-free DMEM before adding 50μCi of L-[35S]-methionine (Perkin Elmer) for a further 30 min. Cells were then washed in PBS and total lysates analysed by SDS-polyacrylamide gel electrophoresis. Following fixation in 50% v/v ethanol and 10% v/v acetic acid, the gel was vacuum dried onto Whatman filter paper and exposed to X-ray film overnight.

### SDS-PAGE and Western blotting

Protein samples were analysed by SDS-polyacrylamide gel electrophoresis and transferred to nitrocellulose membrane for Western blot analysis. Western blots were developed using SuperSignal West Pico chemiluminescent substrate.

### RNA-Seq: Library preparation, sequencing and assembly

Total RNA was extracted from 1 × 10^6^ cells using Qiagen RNeasy reagents, with seven biological replicates prepared for each condition. The quality of the RNA preparations was assessed using a bioanalyser and five biological replicates representing each condition chosen for downstream library preparation. Illumina RNA-Seq sequence libraries were constructed using the Strand-Specific RNA reagent kit (Agilent Technologies, G9691A), according to manufacturer’s instructions (Protocol Version E0, March 2017). Here 1 µg of total RNA was used as input for each sample. Sequence libraries were subsequently QC’d, multiplexed, and run on an Illumina NextSeq 550 (75 cycle, high output) resulting in paired-end (2 × 36 bp) datasets. Following the analysis of this initial dataset which indicated vastly different compositions of the uninfected and 12 hpi infected samples that made it difficult to determine differential expression with confidence, a further set of RNA samples was prepared from uninfected, 12 hpi Wt and 12 hpi Δ22 infected HFFF cells in biological triplicate. Here 1 µg of total RNA was again used as input for each sample but this time was spiked with 2 µL of a 1:100 dilution of the ERCC RNA Spike-In Mix 1 (ThermoFisher, 4456740). Sequence libraries were subsequently QC’d, multiplexed, and run on an Illumina NextSeq 550 (75 cycle, high output) resulting in paired-end (2 × 36 bp) datasets.

### Bioinformatic analysis of RNAseq data

#### Preprocessing

Quality checks were performed via FastQC (version 0.11.4) [86]. Trimmomatic tool (version 0.32) [87] was used for quality trimming and clipping of adapters and repeated sequences. Sequencing reads were mapped to the human transcriptome (iGenome file, Homo sapiens UCSC hg19) using Tophat [88] and to the human herpesvirus 1 strain 17 (JN555585.1) coding sequences using Bowtie2 [89]. The function featureCounts from the R package Rsubread [90] was used to assign mapped sequencing reads to genomic features. Genomic features of the host were defined by the tool’s in-built NCBI RefSeq annotations for the hg19 genome and the R package org.Hs.eg.db [91] was used to annotate the genomic features. Filtering of lowly expressed genes was performed by keeping genes with at least 5 counts per million (CPM) in at least 2 samples.

#### Normalisation

We sequenced three additional samples of each experimental condition, all of which had been spiked with the external RNA control consortium (ERCC) spike-in control mix [66, 67], to determine the relationship between RNA-seq read counts and known inputs. We fit a linear regression on the abundance estimated from RNA-seq and the known ERCCs input amounts to derive a scaling factor. We then used the average of the scaling factor, across replicates, as the global normalisation factor per condition.

#### Differential expression analyses

Based on the R Bioconductor package EdgeR [92], CPM values were fitted to a negative binomial generalised log-linear model (GLM) using empirical Bayes tagwise dispersions to estimate the dispersion parameter for each gene. Differential expression in selected contrasts was identified using GLM likelihood ratio tests.

#### Scatter and volcano plots

In all plots host genes are shown in black and virus genes are shown in green. Scatter plots show the mean log2 (CPM+1), across replicates, in each axis, with the red diagonal broken line indicating no change between experimental conditions. In the volcano plots the x-axis corresponds to the log2 FC between experimental conditions and the y-axis corresponds to BH corrected –log10 p-values, with the red horizontal broken line indicating a BH corrected p-value of 0.05.

### Quantitative RT-PCR (qRT-PCR)

Total RNA was extracted from cells using Qiagen RNeasy kit. Excess DNA was removed by incubation with DNase I (Invitrogen) for 15 min at room temperature, followed by inactivation for 10 min at 65°C in 25 nM of EDTA. Superscript III (Invitrogen) was used to synthesise cDNA using random primers according to manufacturer’s instructions. All qRT-PCR assays were carried out in 96-well plates using MESA Blue qPCR MasterMix Plus for SYBR Assay (Eurogentec). Primers for cellular and viral genes are shown in Table S7. Cycling was carried out in a Lightcycler (Roche), and relative expression was determined using the ΔΔCT method [93], using 18s RNA as reference. For validation experiments, total RNA was spiked commercial luciferase RNA (Promega) and relative expression was normalised to the level of luciferase RNA. Statistical analysis was carried out using an unpaired, two-way student’s t test.

### Fluorescent *in situ* hybridisation (FISH) of mRNA

Cells were grown in 2-well slide chambers (Fisher Scientific) and infected with virus. At the appropriate time, cells were fixed for 20 min in 4% PFA, then dehydrated by sequential 5 min incubations in 50%, 70% and 100% ethanol. FISH was then carried out using Applied Cell Diagnostics (ACD) RNAscope reagents according to manufacturer’s instructions. Briefly, cells were rehydrated by sequential 2 min incubations in 70%, 50% ethanol and PBS, and treated for 30 min at 37 **°**C with DNase, followed by 15 min at room temperature with protease. Cells were then incubated for 2 h at 40 **°**C with the relevant RNAscope probe (ICP27; ICP0; TK; gD; gC; VP16; vhs; serpin E1; MMP1; GLUL; POLR2A; PPIB; IFIT1 as designed by Advanced Cell Dignostics, ACD), followed by washes and amplification stages according to instructions. After incubation with the final fluorescent probe, the cells were mounted in Mowiol containing DAPI to stain nuclei, and images acquired with a Nikon A2 inverted confocal microscope.

### Immunofluorescence

Cells grown on coverslips were treated as described previously [94]. Images were acquired on a Nikon A2 confocal microscope and processed using Adobe Photoshop software.

## Acknowledgments

We thank the UCL Pathogen Genomic Unit for performing sequencing reads.

We also thank Steve Triezenberg, Tony Minson, John Wills, Frazer Rixon, Duncan Wilson and David Johnson for generously providing reagents used in this study.

## Supporting information

**S1 Table. RNAseq data for total Wt infected HFFF cell transcriptome at 4 hours compared to uninfected HFFF cell transcriptome.** Raw counts and counts per million for all cell and virus genes in each biological replicate are listed, with genes expressed at a low level filtered out by keeping genes with at least 5 counts per million (CPM) in at least 2 samples. Genes are ordered according to highest to the lowest Log_2_ FC.

**S2 Table. RNAseq data for total Wt infected HFFF cell transcriptome at 12 hours compared to uninfected HFFF cell transcriptome.** Raw counts and counts per million for all cell and virus genes in each biological replicate are listed, with genes expressed at a low level filtered out by keeping genes with at least 5 counts per million (CPM) in at least 2 samples. Genes are ordered according to highest to the lowest Log_2_ FC.

**S3 Table. RNAseq data for total Δ22 infected HFFF cell transcriptome at 4 hours compared to uninfected HFFF cell transcriptome.** Raw counts and counts per million for all cell and virus genes in each biological replicate are listed, with genes expressed at a low level filtered out by keeping genes with at least 5 counts per million (CPM) in at least 2 samples. Genes are ordered according to highest to the lowest Log_2_ FC.

**S4 Table. RNAseq data for total Δ22 infected HFFF cell transcriptome at 12 hours compared to uninfected HFFF cell transcriptome.** Raw counts and counts per million for all cell and virus genes in each biological replicate are listed, with genes expressed at a low level filtered out by keeping genes with at least 5 counts per million (CPM) in at least 2 samples. Genes are ordered according to highest to the lowest Log_2_ FC.

**S5 Table. RNAseq data for total Δ22 infected HFFF cell transcriptome at 4 hours compared to Wt infected HFFF cell transcriptome at 4 hours.** Raw counts and counts per million for all cell and virus genes in each biological replicate are listed, with genes expressed at a low level filtered out by keeping genes with at least 5 counts per million (CPM) in at least 2 samples. Genes are ordered according to highest to the lowest Log_2_ FC.

**S6 Table. RNAseq data for total Δ22 infected HFFF cell transcriptome at 4 hours compared to Wt infected HFFF cell transcriptome at 4 hours.** Raw counts and counts per million for all cell and virus genes in each biological replicate are listed, with genes expressed at a low level filtered out by keeping genes with at least 5 counts per million (CPM) in at least 2 samples. Genes are ordered according to highest to the lowest Log_2_ FC.

**S7 Table. Primer pair sequences used for qRT-PCR.**

**S1 Fig. Translational shutoff and plaque size phenotype of HSV1 lacking either the UL13 or ICP34.5 gene on HFFF cells.**

**S2 Fig. Expression heatmap of interferon-stimulated genes in HSV1 infected cells at 4 and 12 hours after infection.**

**S3 Fig. Validation of RNAseq data by qRT-PCR. Two replicate RNA samples were subjected to qRT-PCR using primers for the indicated transcripts, and the Log_2_ FC compared to that determined in the RNAseq experiment detailed in S2 Table.**

**S4 Fig. Dual transcriptomic analysis of HFFF cells infected with Δ22 HSV1. Differential expression analysis of cell and virus transcripts was conducted using EdgeR as described in Methods. Differences in the number of reads mapped to cell (black circles) and virus (green circles) transcripts were plotted as scatter plots (left hand panel) and volcano plots (right hand panel) comparing results at 4 and 12 hours to uninfected cells.**

**S5 Fig. Relative expression of virus transcriptome in Wt and Δ22 infected HFFF cells.**

